# Polygenic risk scores of several subtypes of epilepsies in a founder population

**DOI:** 10.1101/728816

**Authors:** Claudia Moreau, Rose-Marie Rébillard, Stefan Wolking, Jacques Michaud, Frédérique Tremblay, Alexandre Girard, Joanie Bouchard, Berge Minassian, Catherine Laprise, Patrick Cossette, Simon Girard

**Author notes:** Corresponding author: Simon Girard.

## Abstract

**Importance:** Epilepsy is defined as a group of neurological disorders characterized by epileptic seizures, brief episodes of symptoms that are caused by abnormal or excessive neuronal activity in the brain. Epilepsy affects around 3 percent of individuals. In the past 10 years, many groups have been working to better understand the complex genetic mechanisms underlying epilepsy. Together, they studied many different genetic mechanisms, but there is still a substantial missing heritability component in epilepsy genetics.

**Objective:** Here, we used polygenic risk scores (PRS) to quantify the cumulative effects of a number of variants, which may individually have a very small effect on susceptibility.

**Design:** We calculated PRS in 522 French-Canadian epilepsy patients divided into seven subtypes and French-Canadian controls.

**Setting:** All study participants (cases and controls) were selected based on their French-Canadian ancestry.

**Participants:** The epilepsy cohort was composed of families of at least three affected individuals with Idiopathic Generalized Epilepsy (IGE) or Non-acquired Focal Epilepsy (NAFE) previously collected and diagnosed by neurologists following the International League Against Epilepsy (ILAE) criteria.

**Exposures:** All samples were processed on a common genotyping array.

**Main outcomes and Results:** We show that the area under the curve (AUC) is almost always slightly greater than 0.5, especially in patients with IGE and subtypes. We also looked at the association of the PRS with the different phenotypes using a linear mixed effects model estimated by generalized estimating equation (GEE) with the pairwise identity-by-descent (IBD) matrix as a random effect. P-values of GEE were consistent with AUC calculations.

**Conclusions and Relevance:** Globally, we support the notion that PRS and SNP-based heritability provide reliable measures to rightfully estimate the contribution of genetic factors to the pathophysiological mechanism of epilepsies, but further studies are needed on PRS before they can be used clinically.

## INTRODUCTION

Epilepsy is defined as a group of neurological disorders characterized by epileptic seizures or brief episodes of symptoms that are caused by abnormal or excessive neuronal activity in the brain ^1,2^. Epilepsy affects ∼3% of individuals, with half of these cases starting during childhood. Although monogenic forms of the disease have been reported (with genes such as GABRA1 ^3^, SCN1A ^4^, CHRNA4 ^5^, and LGI1 ^6^), they represent less than 2% of epilepsy cases. In the past 10 years, many groups have been working to better understand the complex genetic mechanisms underlying epilepsy ^7^. Together, they studied many different genetic mechanisms, from monogenic to linkage and copy number variant (CNV) studies, and finally big genome-wide association studies (GWAS) were made possible by next-generation sequencing. The utility of using epilepsy sub-phenotypes has been demonstrated in juvenile myoclonic epilepsy (JME) ^3,8^ but not in other sub-phenotypes. Last year, a large GWAS on epilepsy identified 16 loci associated with the disease, and many of these were already known or suspected ^9^. Despite these efforts, there is still a substantial missing heritability component in epilepsy genetics ^10^.

The heritability calculations in epilepsy greatly differ depending on the method used and the patients analyzed. In a recent study ^9^, 32% single-nucleotide polymorphism (SNP) heritability for Idiopathic Generalized Epilepsy (IGE) and only 9% for Non-acquired Focal Epilepsy (NAFE) was calculated. In another study ^11^, 26% heritability on common single nucleotide polymorphisms (SNPs) (>0.01) was calculated for all epilepsies and 27% heritability for NAFE patients. This again denotes the heterogeneous genetic landscape of the disease, the variety of methods used, and also the need to study the different epilepsy sub-phenotypes.

It is likely that a wide spectrum of genetic factors is in play, ranging from very rare mutations with large effects to relatively rare variants with medium effect sizes, and finally to common variants with smaller risk effects. Polygenic risk scores (PRS) aim to quantify the cumulative effects of a number of variants, which may individually have a very small effect on susceptibility. They can be used to estimate a person’s likelihood of displaying any trait with a genetic component. They have been used previously in many common traits and diseases such as heart disease ^12–14^ and more importantly in neurological disorders such as schizophrenia 1^5–18^.

In the present study, we take advantage of the most recent meta-analysis GWAS metrics^9^ to calculate PRS in 522 French-Canadian epilepsy patients divided into seven subtypes. The French-Canadian population is known for its well documented recent (400 years) founder effect and its particular genetic background, which makes it an ideal population for genetic studies. French-Canadians are also closely related to the European population, which is predominant in the International League Against Epilepsy (ILAE) study. The French-Canadian population has been successfully used to confirm the ability of PRS to predict the risk of another polygenic disease, coronary artery disease that was initially conducted in Europeans ^13^.

In this study, we show that the French-Canadian epilepsy patients share a significant fraction of SNP-based heritability with that reported in the ILAE study, and we also report some of the first evidence that PRS can be used to positively discriminate between cases and controls for several types of epilepsies.

## DATA AND METHODS

### Phenotyping of patients

This study was approved by the CHUM ethics committee and all participants signed a consent form. The epilepsy cohort was composed of families of at least three affected individuals with IGE or NAFE previously collected and diagnosed by neurologists. The clinical epilepsy phenotype is defined based on the Classification of the Epilepsy Syndromes established by the International League against Epilepsy (ILAE) ^19^. More specifically, the operational definitions of the epilepsy phenotypes studied in the project for NAFE were: 1) Patients were at least 5 years of age and had experienced at least two unprovoked seizures in the 6 months prior to starting treatment AND 2) an MRI scan of the brain that did not demonstrate any potentially epileptogenic lesion (No Lesion) OR 3) documented hippocampal sclerosis (HS) and lesion other than mesial temporal sclerosis (Other Lesion).

For IGE, the operational definitions were: 1) Patients with clinical and electroencephalogram characteristics meeting the 1989 ILAE syndrome definitions for childhood absence epilepsy (CAE), juvenile absence epilepsy (JAE), juvenile myoclonic epilepsy (JME), or IGE not otherwise specified 2) All patients were at least 4 years of age at the time of diagnosis. In IGE, we also included patients with epilepsy with eyelid myoclonia (Jeavons), which is an idiopathic generalized form of reflex epilepsy characterized by childhood onset, unique seizure manifestations, striking light sensitivity, and possible occurrence of generalized tonic-clonic seizures alone (GTCS).

Our FC epilepsy cohort consisted of 643 patients diagnosed with epilepsy. French Canadian (FC) ancestry was assessed by self-declared ethnicity and principal component analysis (eFigure 1). One hundred twenty-one individuals were removed based on ethnicity, leaving a cohort of 522 FC epilepsy patients of which 262 patients had IGE and 163 patients were diagnosed with a NAFE. Table 1 shows the different subtypes of epilepsies that are represented in our cohort. Additionally, we selected 954 French-Canadian individuals from a reference population ^20^.

**Table 1:** Cohorts description.

| **Epilepsy subtype** | **n** | **n women** | **Mean age** |
| --- | --- | --- | --- |
| All epilepsies | 522 | 300 | 48 |
| IGE (including all syndromes and unspecified) | 262 | 154 | 45 |
| CAE | 43 | 21 | 36 |
| JAE | 28 | 17 | 50 |
| JME | 102 | 67 | 47 |
| GTCS | 32 | 20 | 42 |
| NAFE (including all types and unspecified) | 163 | 83 | 49 |
| NAFE HS | 25 | 11 | 52 |
| NAFE no lesion | 81 | 45 | 45 |
| NAFE other lesion | 16 | 11 | 54 |

\*Types of epilepsies : IGE = Idiopathic Generalized Epilepsy, CAE = Childhood Absence Epilepsy, JAE = Juvenile Absence Epilepsy, JME = Juvenile Myoclonic Epilepsy, GTCS = generalized tonic-clonic seizures alone, NAFE = Non-Acquired Focal Epilepsy, HS =
documented hippocampal sclerosis, No lesion: no documented epileptogenic lesion, Other lesion: lesion other than mesial temporal sclerosis.

**Figure 1:**
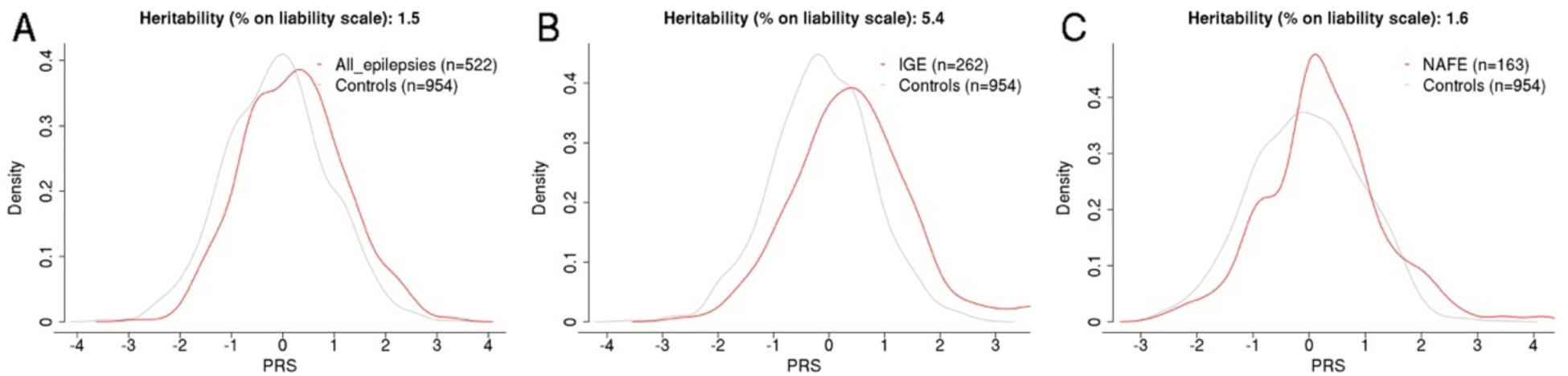
PRS density plots and heritability on liability scale for A) all epileptic patients, B) IGE patients and C) NAFE patients.

### Genotyping and imputation

For this study, we used whole-genome genotyping data for the patient and the French-Canadian control ^20^ cohorts. All samples were processed on either the Illumina Omni Express (n.SNPs = 710,000) or the Illumina Omni 2.5 (n.SNPs = 2,500,000 including the Omni Express core). Genotypes of all samples were merged and only positions present on both chips were kept. We performed cleaning steps to remove individuals having more than 2% missing genotypes among all SNPs, SNPs with more than 2% missing SNPs over all individuals, and SNPs with HWE p-value < 0.001 using PLINK software ^21^. We then removed 121 individuals of non-FC descent using the first two principal components (PCs) in addition to self-identification of patients whenever this information was available. PCA was performed using Eigensoft ^22^ on pruned SNPs (pairwise r^2^ < 0.2 in sliding windows of size 50 shifting every five SNPs) at 5% minor allele frequency (MAF). We finally aligned the dataset to the GRCh37 genome build for further imputation following the method described here ^23^.

The Sanger Imputation Service was used to conduct whole-genome imputation of SNPs ^24^. We selected the Human Reference Consortium dataset as the reference panel. Post-imputation quality control filters were applied to remove SNPs within imputed data with an imputation info score <0.9 or HWE p-value <1e-6, and only biallelic SNPs at MAF 1% or higher were kept for further analyses.

### PRS calculation

PRS were calculated with PRSice software ^25^ using ILAE meta-analysis on epilepsy summary statistics ^9^. Since the BETA was not provided for the METAL analyses (all, generalized, and focal epilepsies’ analyses), we used a fixed BETA of 0.001 or −0.001 depending on the z-score direction for these PRS calculations. We used the first 10 PCs in addition to sex as covariates, recalculating eigenvectors for each patient subset including controls using SNPs at MAF 0.05 pruned (as described above). PRS were standardized for graphs.

### Statistical analyses

We used PLINK software for the GWAS analysis with the first 10 PCs as covariates. We used the Genetic Analysis Repository software ^26^ to estimate heritability (i.e., variance explained at the liability scale) with the Nagelkerke’s pseudo-R^2^ (using the PRS at the p-value threshold that best predicts the phenotype) assuming a liability-threshold model. a prevalence of 0.002 and 0.003 for IGE and NAFE, respectively (and 0.002 for all cases together), and adjusting for case-control ascertainment.

To examine the ability of PRS to predict the epilepsy phenotype, we calculated the area under the curve (AUC) using the pROC package in R. We also used the lmekin function of the coxme R package that fits a linear mixed effects model estimated by generalized estimating equations (GEE) with random effects, which is robust for intra-familial correlation. We used the identity-by-descent (IBD) correlation matrix calculated between each pair of individuals using PLINK.

## RESULTS

We used whole-genome genotyping on 522 patients suffering from epilepsy and 954 population controls. All individuals were confirmed with French-Canadian ancestry. The first step of our study was to assess which of the associations found by the ILAE study ^9^ were valid for our cohort. Table 2 presents the GWAS-associated SNPs (only the lead SNP was tested) as well as the association statistics for our cohort. Associations were only tested in the epilepsy subtypes in which they were originally reported. When correcting for multiple testing using the Bonferroni correction (n = 20), three loci were found to be significant, all within the IGE epilepsy subtype. These results show that our founder FC population shares a significant portion of the epilepsy genetic risks with the populations studied by the ILAE.

**Table 2:** Replicated GWAS hits.

| Epilepsy subtype | SNP | Gene(s) | MAF patients | MAF controls | OR | Chi<sup>2</sup> | P-value |
| --- | --- | --- | --- | --- | --- | --- | --- |
| All epilepsies | rs4671319 | FANCL,BCL11A | 0.49 | 0.45 | 1.22 | 2.42 | 0.016 |
|  | rs6432877 | SCN3A,SCN2A,TTC21B,SCN1A | 0.28 | 0.27 | 1.03 | 0.27 | 0.789 |
|  | rs4638568 | HEATR3,,BRD7 | 0.08 | 0.08 | 0.92 | -0.51 | 0.611 |
| IGE | **rs4665630** | **None** | **0.16** | **0.12** | **1.66** | **3.18** | **0.001** |
|  | **rs1402398** | **FANCL,BCL11A** | **0.48** | **0.37** | **1.71** | **4.73** | **0.000002** |
|  | rs11890028 | SCN3A,SCN2A,TTC21B,SCN1A | 0.30 | 0.30 | 1.01 | 0.07 | 0.944 |
|  | rs887696 | STAT4 | 0.39 | 0.35 | 1.19 | 1.49 | 0.135 |
|  | rs1044352 | PCDH7 | 0.42 | 0.39 | 1.09 | 0.72 | 0.473 |
|  | rs11943905 | GABRA2 | 0.32 | 0.30 | 0.99 | -0.12 | 0.902 |
|  | rs4596374 | KCNN2 | 0.45 | 0.43 | 1.05 | 0.39 | 0.698 |
|  | rs68082256 | ATXN1 | 0.21 | 0.23 | 0.92 | -0.62 | 0.538 |
|  | rs13200150 | None | 0.26 | 0.31 | 0.81 | -1.70 | 0.090 |
|  | rs4794333 | PNPO | 0.36 | 0.37 | 0.89 | -1.00 | 0.316 |
|  | **rs2833098** | **GRIK1** | **0.28** | **0.38** | **0.64** | **-3.51** | **0.0005** |
| CAE | rs12185644 | FANCL,BCL11A | 0.43 | 0.31 | 1.81 | 2.46 | 0.014 |
|  | rs13020210 | ZEB2 | 0.20 | 0.15 | 1.32 | 0.94 | 0.350 |
| JME | rs1046276 | STX1B | 0.37 | 0.32 | 1.65 | 2.92 | 0.003 |
| NAFE | rs2212656 | SCN3A,SCN2A,TTC21B,SCN1A | 0.26 | 0.27 | 0.82 | -1.22 | 0.222 |
| NAFE HS | rs1991545 | C3orf33,SLC33A1,KCNAB1 | 0.02 | 0.02 | 1.05 | 0.05 | 0.961 |
|  | rs1318322 | GJA1 | 0.14 | 0.15 | 0.82 | -0.49 | 0.624 |

Next, to assess how SNPs taken together (and not one at a time, like in the GWAS) could explain the missing heritability of epilepsy, we used the basic statistics of the ILAE study^9^ to construct PRS. Figure 1 shows the density plots of standardized PRS values of patients compared to controls for the three broad epilepsy types (for the best-fit p-value, see eFigures 2-4). The PRS for these phenotypes was calculated using a fixed BETA (see methods). Heritability calculated using the Nagelkerke’s pseudo-R^2^ on the liability scale is also shown on each panel. Our first observation was that the PRS distribution is always more shifted to the right and heritability is higher in IGE than in NAFE, which was also reported in this ILEA study ^9^. Figure 2 shows the same analysis for the IGE subtypes CAE (A), GTCS (B), JME (C), and JAE (D) and Figure 3 for the NAFE subtypes HS (A), no lesion (B), or lesions other than HS (C). The best fits are shown in eFigures 5 to S11. Heritability on liability scale varied across the different subtypes; however, it is generally higher for the IGE (3 to 23% and 5.6% for all IGE) types than for the NAFE subtypes, except for the NAFE with lesions other than HS (that has a very low sample size and the lowest best-fit p-value (in total only 6 SNPs were used with a p-value <10^-7^), see eFigure 11).

**Figure 2:**
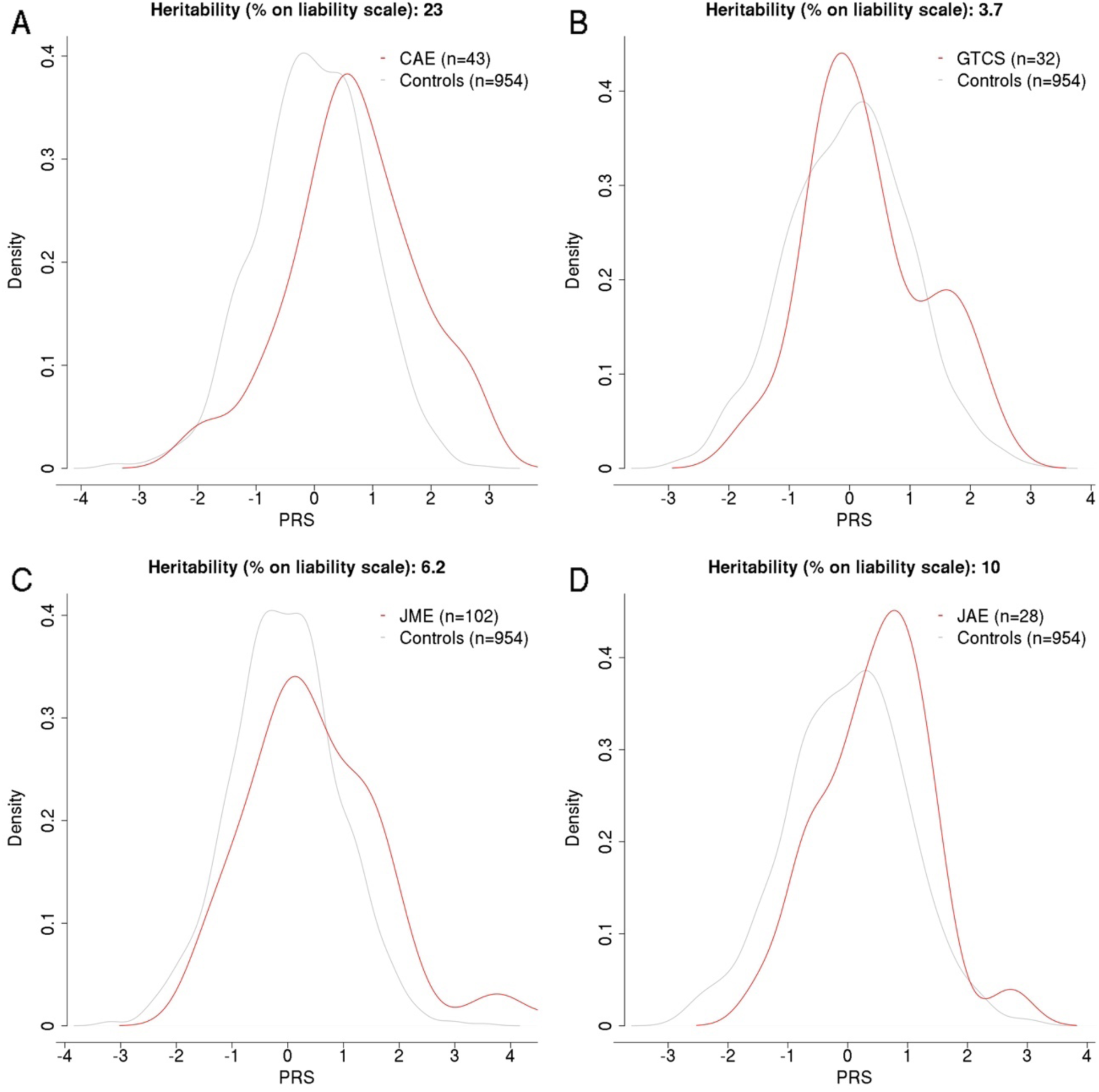
PRS density plots and heritability on liability scale for IGE syndrome A) CAE, B) GTCS, C) JME and D) JAE patients.

**Figure 3:**
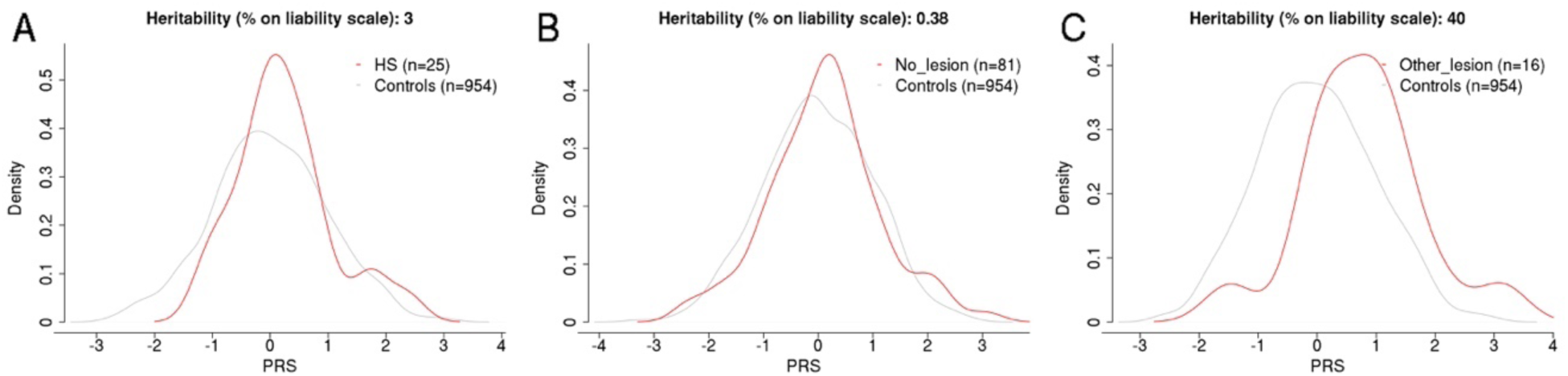
PRS density plots and heritability on liability scale for NAFE patients A) hippocampal sclerosis (HS), B) no documented lesion and C) lesions other than HS.

The next logical question was to investigate if genetic variants taken all together in PRS can be used to discriminate between an epilepsy patient and a control. Table 3 presents the area under the curve (AUC) of the PRS values for patients and controls for all epilepsy subtypes. The ability of PRS to predict the phenotype varies among epilepsy subtypes, but is higher for IGE types (ranging between 0.572 to 0.698) than for NAFE types (ranging between 0.504 to 0.556, excluding the NAFE with other lesions), which was also reflected by the generally higher heritability explained by the PRS in IGE compared to NAFE (see Figures 1–3). To determine the association of the PRS with the phenotype, we performed a linear mixed effects model estimated by GEE with the pairwise IBD matrix as a random effect. Table 3 shows the p-values for the model. Most of the IGE epilepsy subtypes have significant p-values, except GTCS. NAFE patients have less significant effects, and only NAFE with lesion other than HS are significant. However, as stated previously, the calculation of PRS at the best-fit p-value for this subtype included only six SNPs in 16 patients.

**Table 3:** AUC of PRS and GEE p-value for patients and controls for different epilepsy subtypes

| **Epilepsy subtype** | **AUC** | **AUC CI (95%)** | **GEE p-value** |
| --- | --- | --- | --- |
| All epilepsies | 0.59 | 0.5598-0.62 | 1.7E-09 |
| IGE | 0.654 | 0.6156-0.6928 | 0.0E+00 |
| CAE | 0.698 | 0.6109-0.7841 | 2.4E-09 |
| JAE | 0.641 | 0.5413-0.741 | 9.3E-03 |
| JME | 0.624 | 0.5631-0.6845 | 8.2E-08 |
| GTCS | 0.572 | 0.4737-0.6694 | 9.5E-02 |
| NAFE | 0.556 | 0.5063-0.6062 | 9.4E-03 |
| NAFE HS | 0.504 | 0.3781-0.6299 | 9.5E-01 |
| NAFE no lesion | 0.53 | 0.466-0.5935 | 1.0E-01 |
| NAFE other lesion | 0.696 | 0.5729-0.8199 | 2.9E-03 |

## DISCUSSION

In the current study, we confirm the multifactorial nature of epilepsies by replicating several associations identified in the largest GWAS to date conducted on epilepsies. The strongest association in our FC cohort was observed with the SNP rs1402398. This SNP is located in the non-coding region surrounding genes *FANCL* and *BCL11A*. These genes have been linked with epilepsies through association studies ^27^ but no other functional or clinical evidence highlight their roles in the disease. The second replicated association was near a SNP surrounding the *GRIK1* gene. Although this gene encodes a GluR5 kainate receptor gene and its function is plausible in epilepsies, we have limited evidence to decipher its role in the disease ^28^. Finally, the third association was made with SNP rs4665630 found in a genomic region that does not contain any epilepsy-associated genes. Although we successfully replicated several associations, we believe that the biggest contribution of our study lies in the PRS that were established for each epilepsy type. We have shown that the PRS are higher for all epilepsy subtypes when compared to the control cohort. Additionally, we showed that the AUC measure can be used to predict the status of an individual with regards to epilepsy, to some extent. This is particularly true for IGE cases and for several IGE subtypes, such as CAE and JME. This is, to our knowledge, one of the first documented examples of how PRS can be used for epilepsy genetic studies. Although this measure cannot yet be translated to clinical use, our analysis shows that the additive value of common variants can be used to better understand the disease.

One definite pitfall of our study is the small size of our cohort. The initial GWAS was done on more than 15,000 epilepsy patients. Our study only included 522 epilepsy patients and thus cannot have the same outreach as the initial one. This is why we did not report genome-wide association statistics and focused only on the replication of associated SNPs. We believe that the small size of our cohort also impacts the PRS calculations, but to a smaller degree. For instance, the AUC of the PRS and all other statistics for the NAFE without lesion type should be cautiously considered as the cohort size is small (n = 16).

For these reasons, we have to take the heritability explained by the PRS with caution. However, for the broad phenotypes, we explain 5.4% of the heritability for IGE patients, which is higher than what we explain for NAFE patients (1.6%), as expected. This also supports the fact that epilepsy should be divided into subtypes when studying the genetic mechanism underlying the disease, as some epilepsy types were reasonably well-explained by the PRS (i.e., CAE with 23%).

This study was conducted on a documented founder population. The FC population is well-known for its high prevalence of specific disease-causing mutations ^29,30^. For epilepsy, although we cannot exclude that some of the associations found were driven by rare haplotypes, we show here that the genetic etiology of the disease is consistent with that of the general European population. In future work, we will try to assess if the strong PRS found in CAE and JAE could be explained by rarer haplotypes, as we would expect in a founder population.

Globally, we support the notion that PRS and SNP-based heritability provide reliable measures to rightfully estimate the contribution of genetic factors to the pathophysiological mechanism of epilepsies.

## Supporting information

eFigures

## ACKNOWLEDGEMENTS

We are thankful to Compute Canada/Calcul Québec for the access to storage and computing resources. Thanks to Alexandre Bureau for his useful expertise in biostatistics. We would like to thank Editage (www.editage.com) for English language editing. We are extremely grateful to all patients and their families for accepting to participate in this research. We would like to thank Damian Labuda and Hélène Vézina for their work on the QRS cohort. This work was supported by funding from Genome Quebec / Genome Canada.

