## Supplementary material for "Polygenic risk scores of several subtypes of epilepsies in a founder population": eFigures

Online supplement for manuscript *Polygenic risk scores of several subtypes of epilepsies in a founder population*

**In this document:**

**eFigure 1:** PCA of 522 French-Canadian patients with all epilepsy subtypes and 954 French-Canadian controls.
**eFigure 2:** Bar plot from PRSice showing results at broad *P*-value thresholds for epilepsy PRS predicting status for all 522 epileptic patients and 954 French-Canadian controls.
**eFigure 3:** Bar plot from PRSice showing results at broad *P*-value thresholds for epilepsy PRS predicting status for 262 IGE patients and 954 French-Canadian controls.
**eFigure 4:** Bar plot from PRSice showing results at broad *P*-value thresholds for epilepsy PRS predicting status for 163 NAFE patients and 954 French-Canadian controls.
**eFigure 5:** Bar plot from PRSice showing results at broad *P*-value thresholds for epilepsy PRS predicting status for 43 CAE patients and 954 French-Canadian controls.
**eFigure 6:** Bar plot from PRSice showing results at broad *P*-value thresholds for epilepsy PRS predicting status for 32 GTCS patients and 954 French-Canadian controls.
**eFigure 7:** Bar plot from PRSice showing results at broad *P*-value thresholds for epilepsy PRS predicting status for 102 JME patients and 954 French-Canadian controls.
**eFigure 8:** Bar plot from PRSice showing results at broad *P*-value thresholds for epilepsy PRS predicting status for 28 JAE patients and 954 French-Canadian controls.
**eFigure 9:** Bar plot from PRSice showing results at broad *P*-value thresholds for epilepsy PRS predicting status for 25 NAFE HS patients and 954 French-Canadian controls.
**eFigure 10:** Bar plot from PRSice showing results at broad *P*-value thresholds for epilepsy PRS predicting status for 81 NAFE with no documented lesion patients and 954 French-Canadian controls.
**eFigure 11:** Bar plot from PRSice showing results at broad *P*-value thresholds for epilepsy PRS predicting status for 16 NAFE with documented lesion other than HS patients and 954 French-Canadian controls.

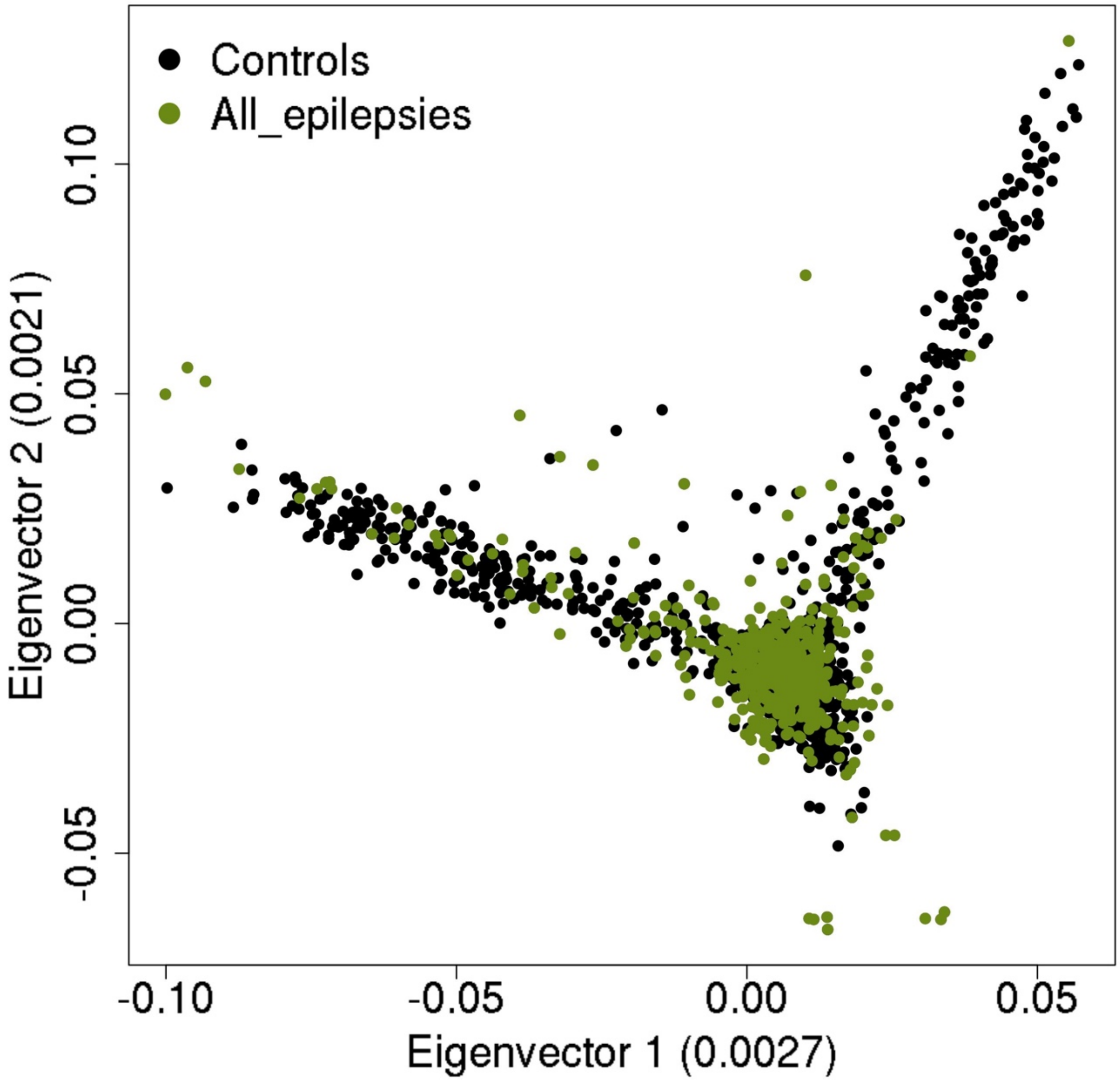


**eFigure 1**: **PCA of 522 French-Canadian patients with all epilepsy subtypes and 954 French-Canadian controls.**

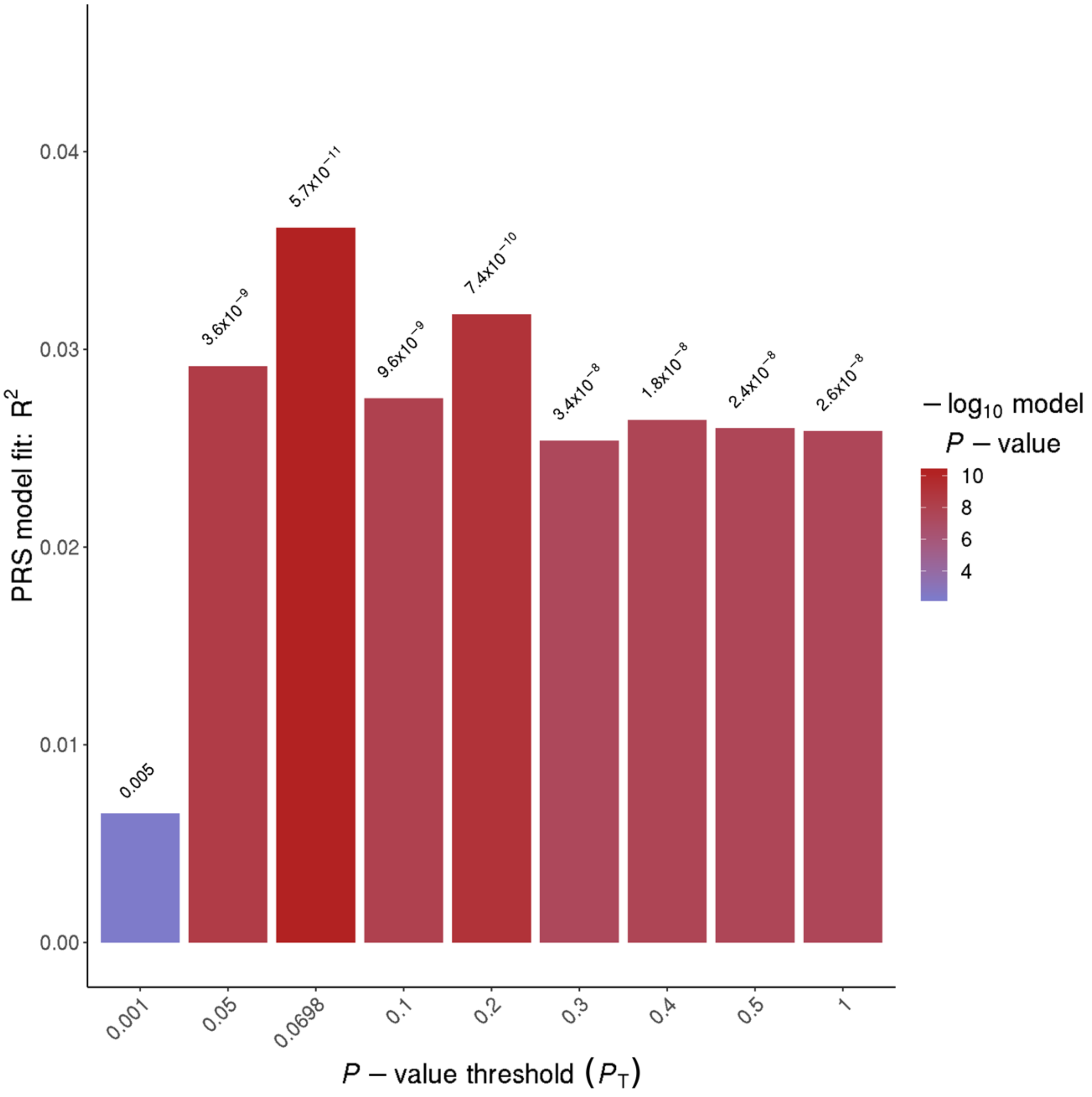


**eFigure 2: Bar plot from PRSice showing results at broad *P*-value thresholds for epilepsy PRS predicting status for all 522 epileptic patients and 954 French-Canadian controls.**

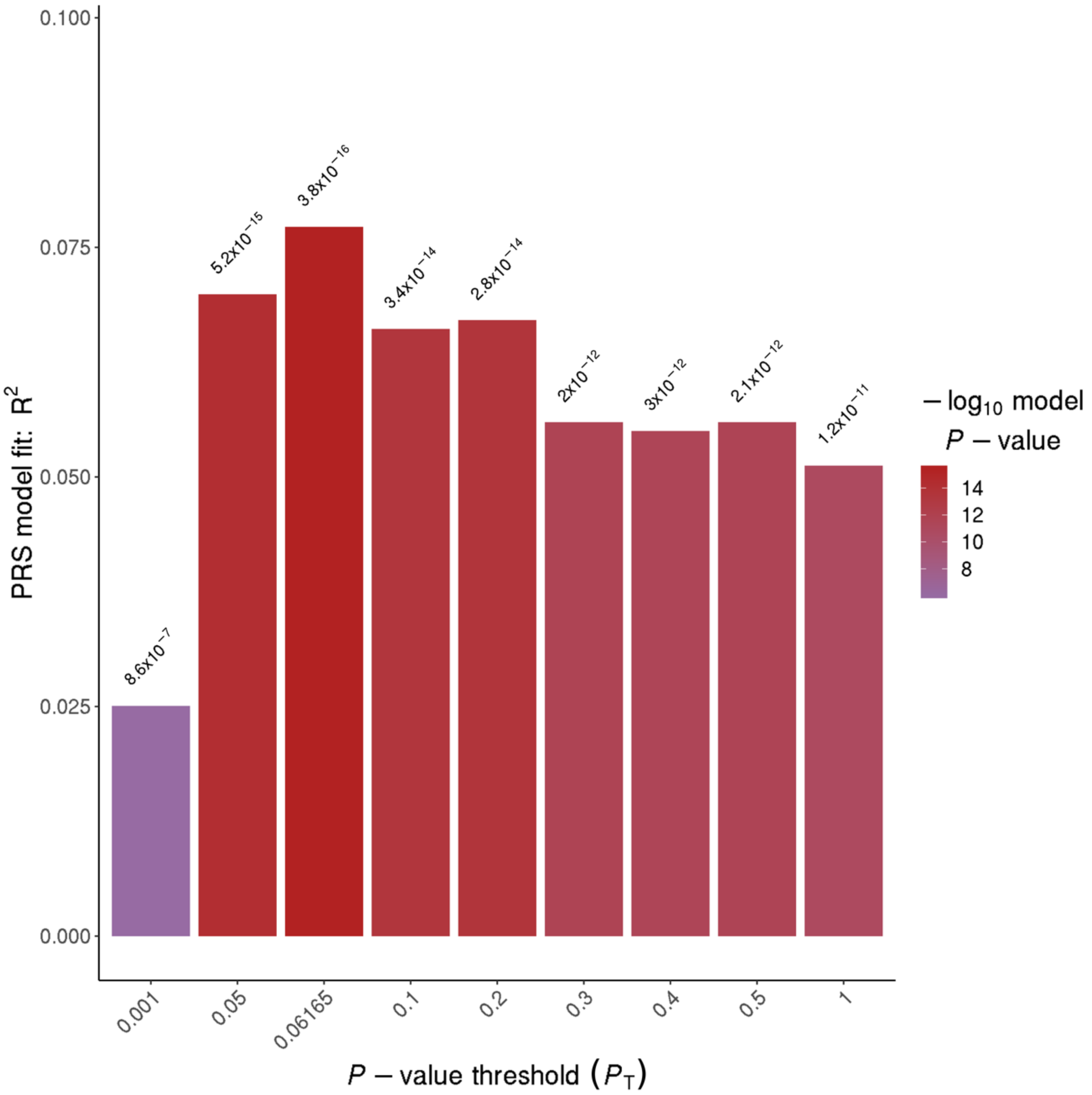


**eFigure 3: Bar plot from PRSice showing results at broad *P*-value thresholds for epilepsy PRS predicting status for 262 IGE patients and 954 French-Canadian controls.**

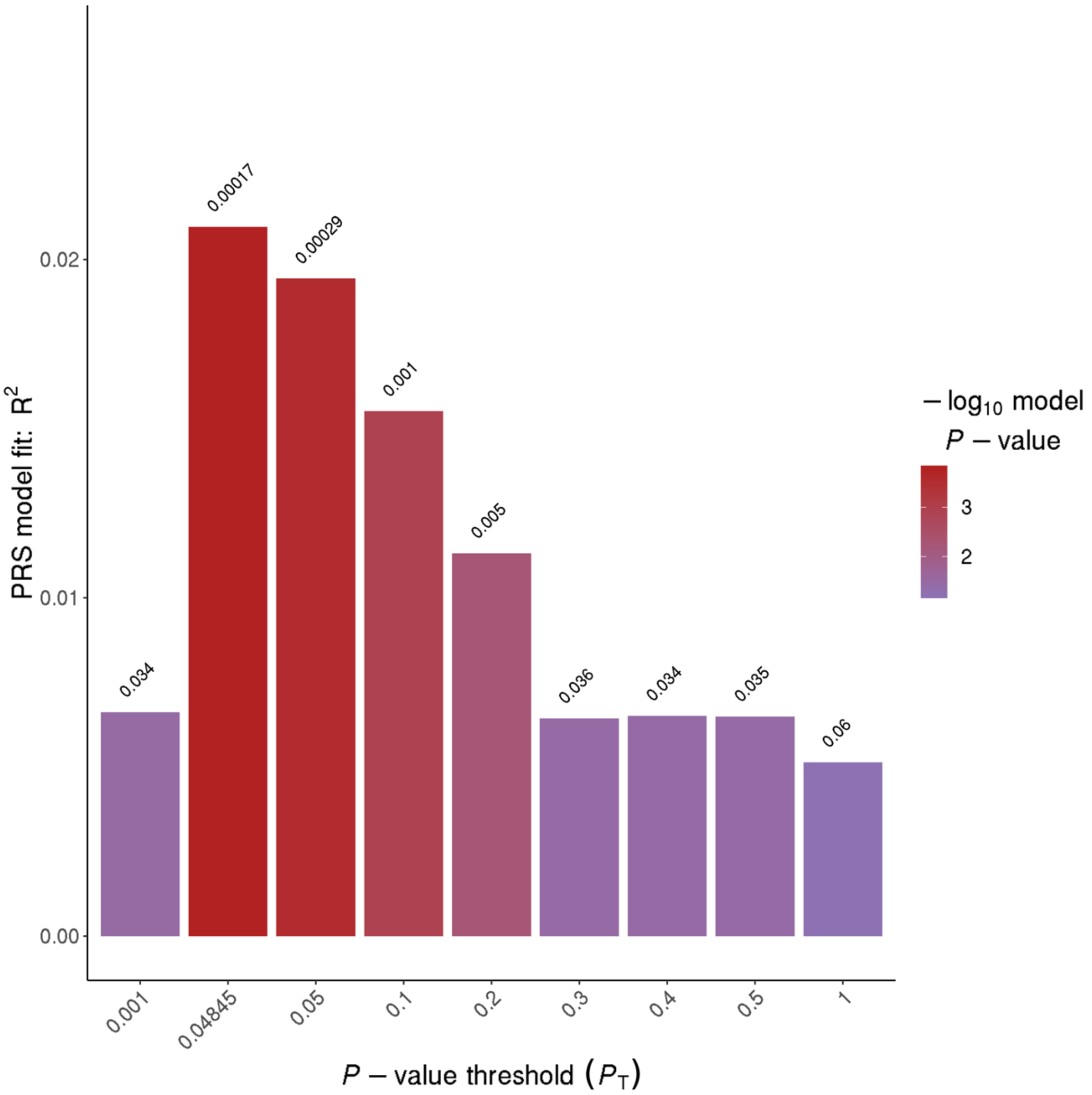


**eFigure 4: Bar plot from PRSice showing results at broad *P*-value thresholds for epilepsy PRS predicting status for 163 NAFE patients and 954 French-Canadian controls.**

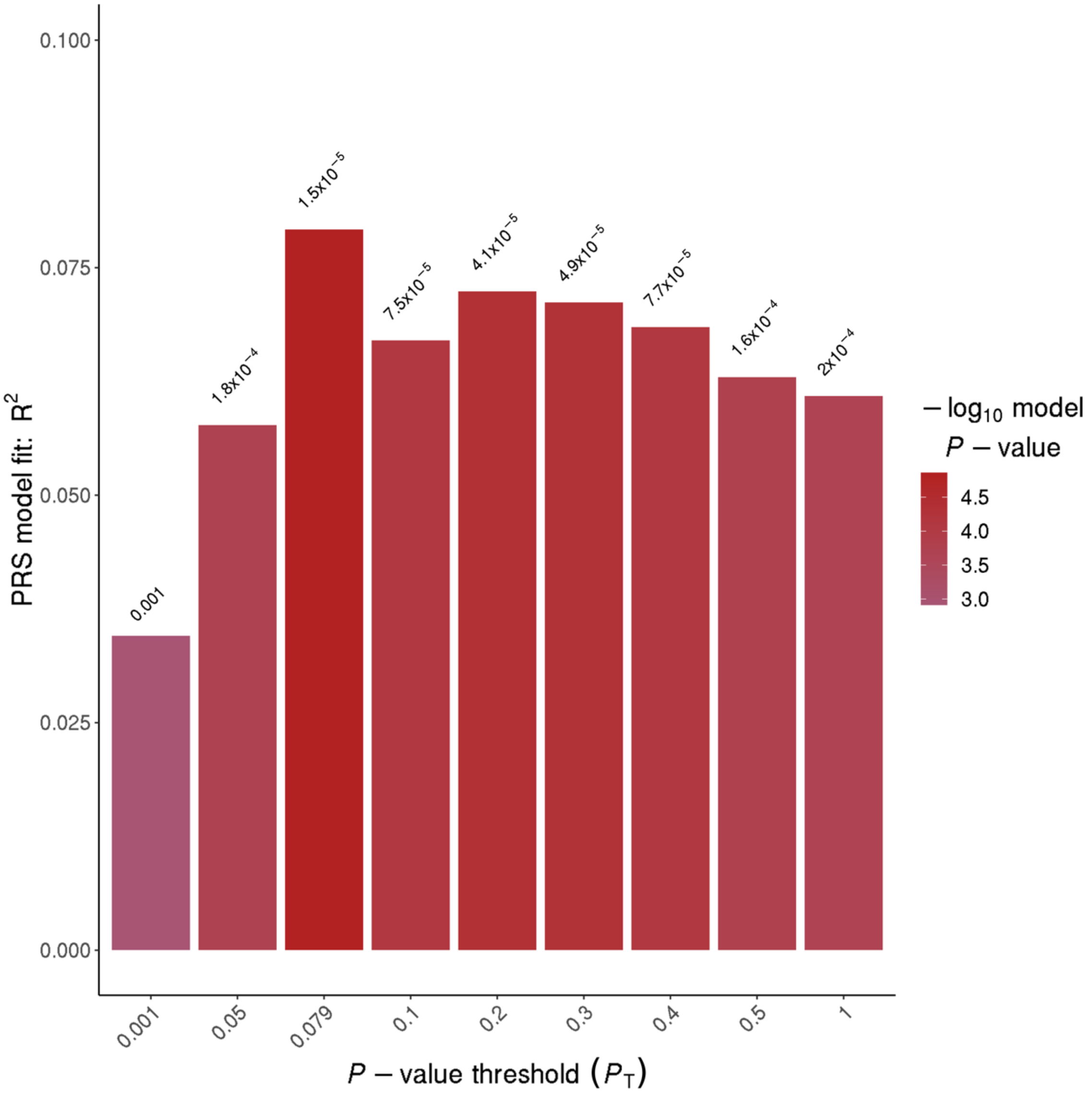


**eFigure 5: Bar plot from PRSice showing results at broad *P*-value thresholds for epilepsy PRS predicting status for 43 CAE patients and 954 French-Canadian controls.**

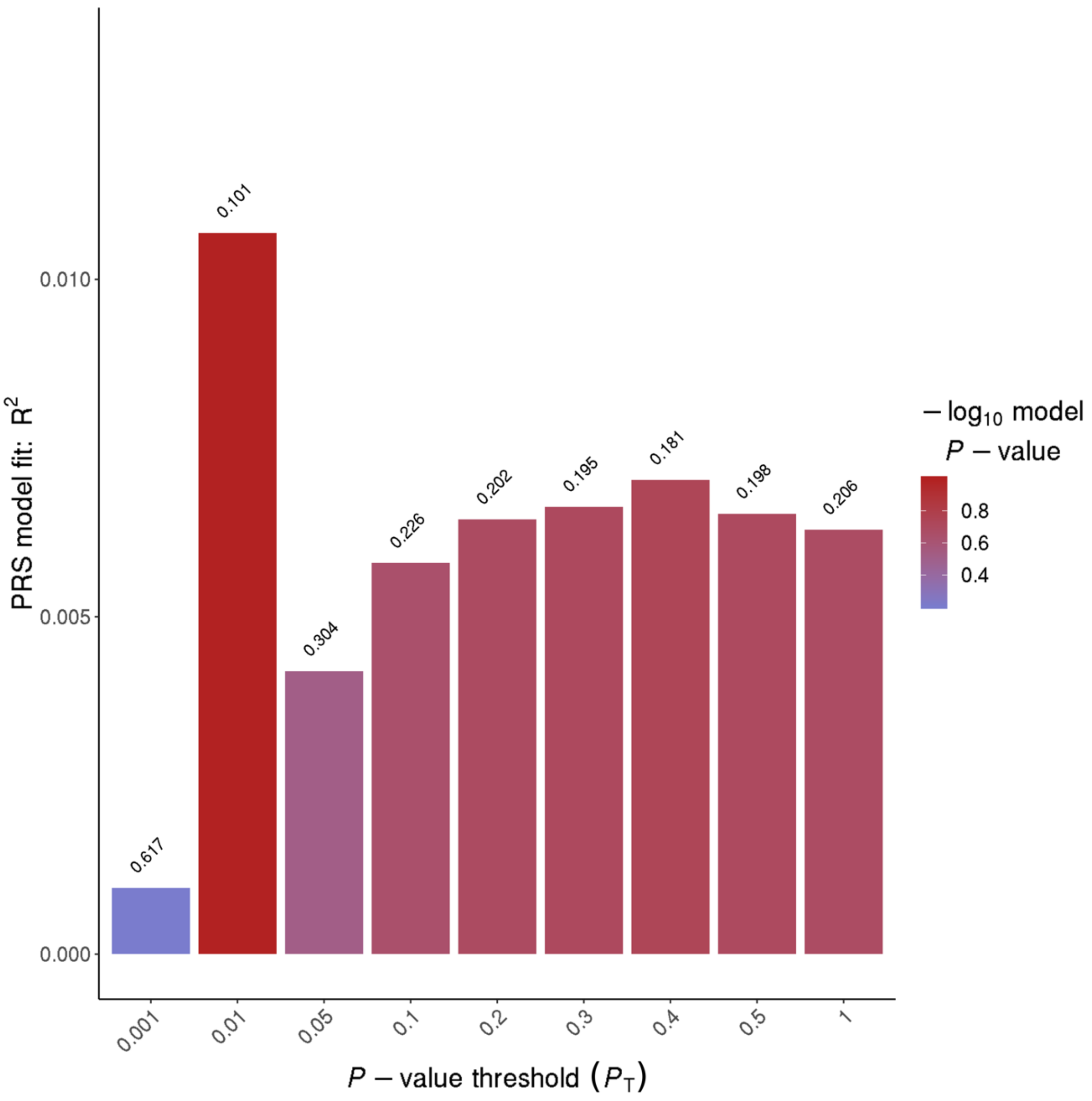


**eFigure 6: Bar plot from PRSice showing results at broad *P*-value thresholds for epilepsy PRS predicting status for 32 GTCS patients and 954 French-Canadian controls.**

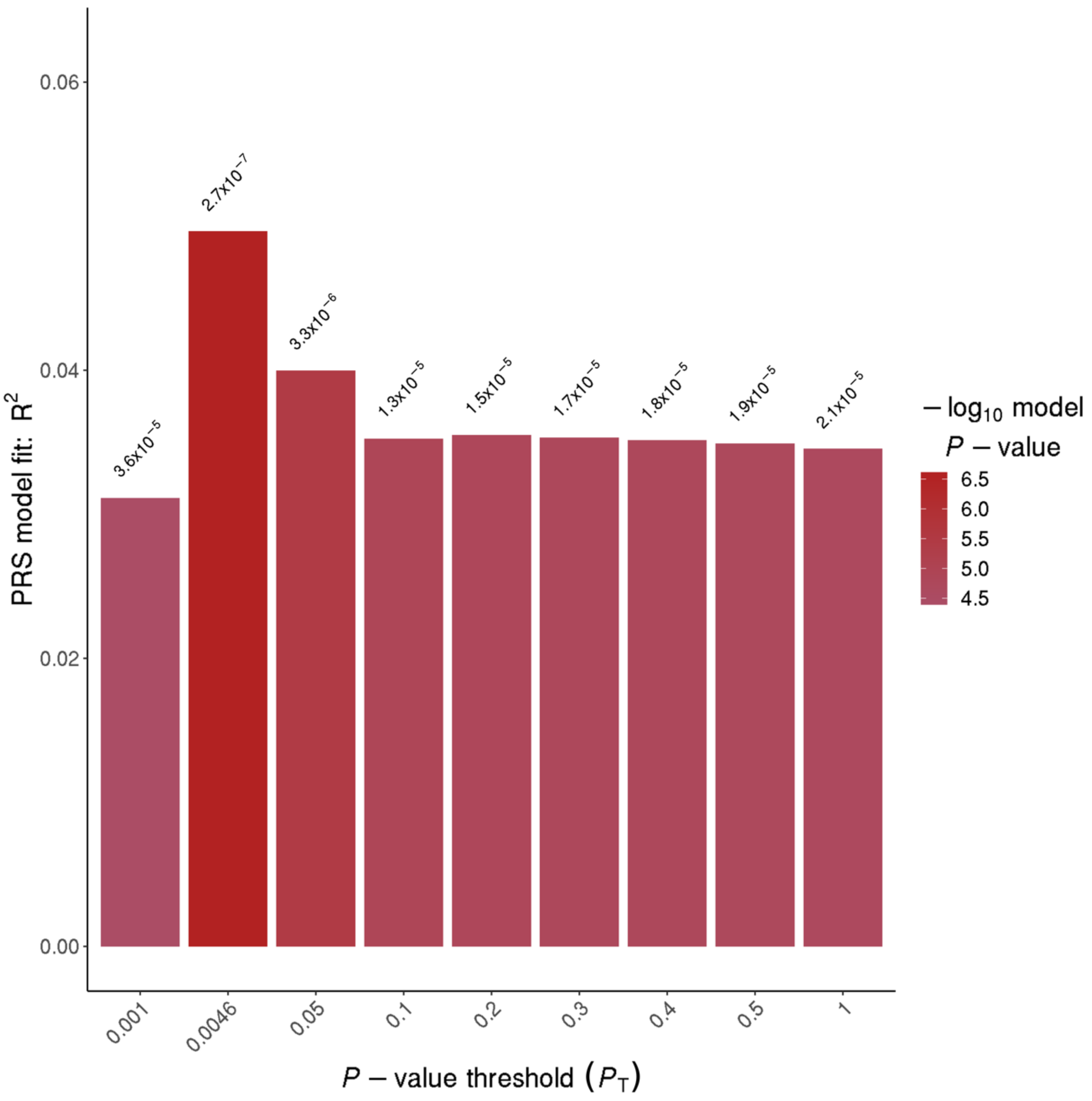


**eFigure 7: Bar plot from PRSice showing results at broad *P*-value thresholds for epilepsy PRS predicting status for 102 JME patients and 954 French-Canadian controls.**

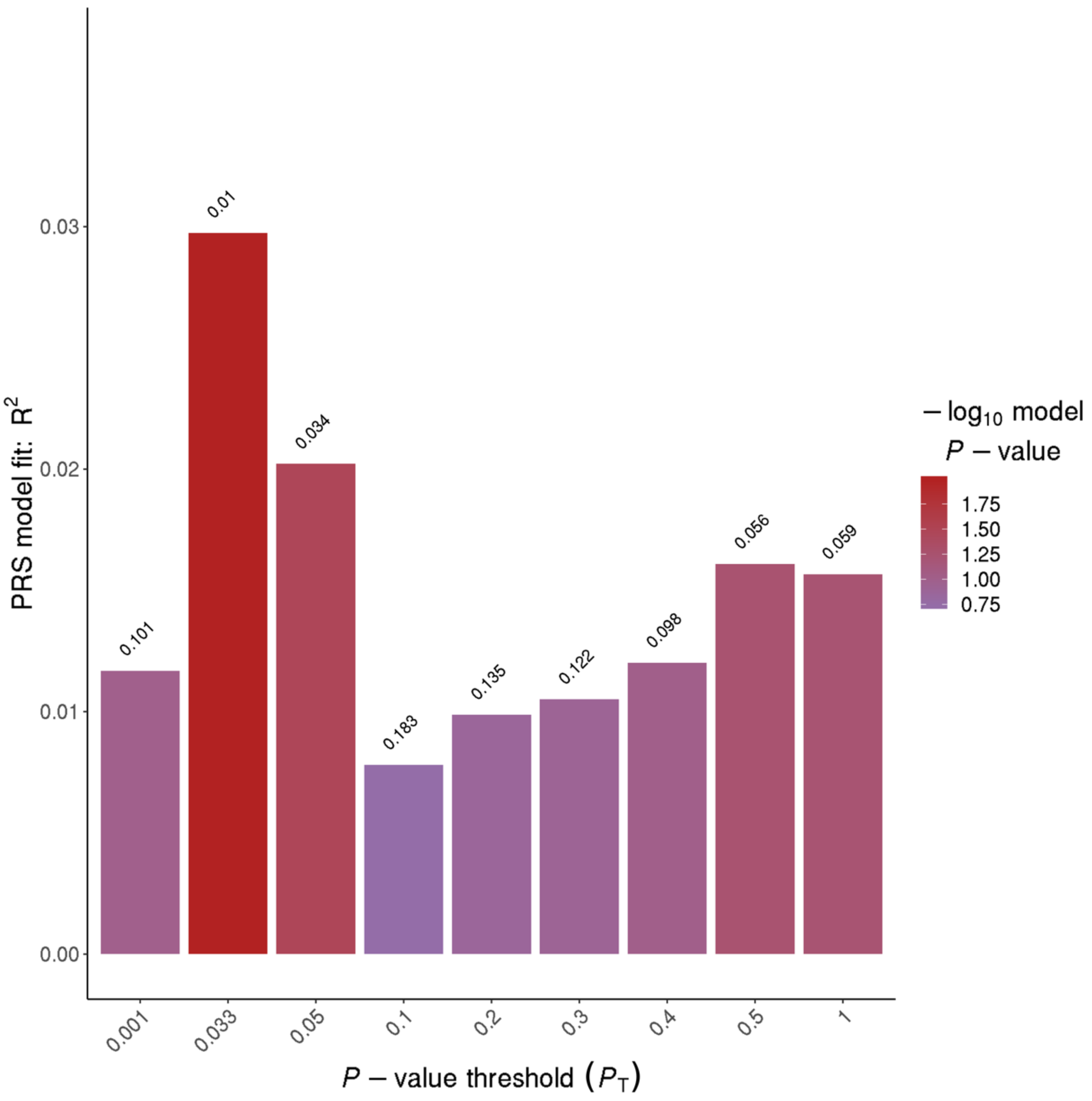


**eFigure 8: Bar plot from PRSice showing results at broad *P*-value thresholds for epilepsy PRS predicting status for 28 JAE patients and 954 French-Canadian controls.**

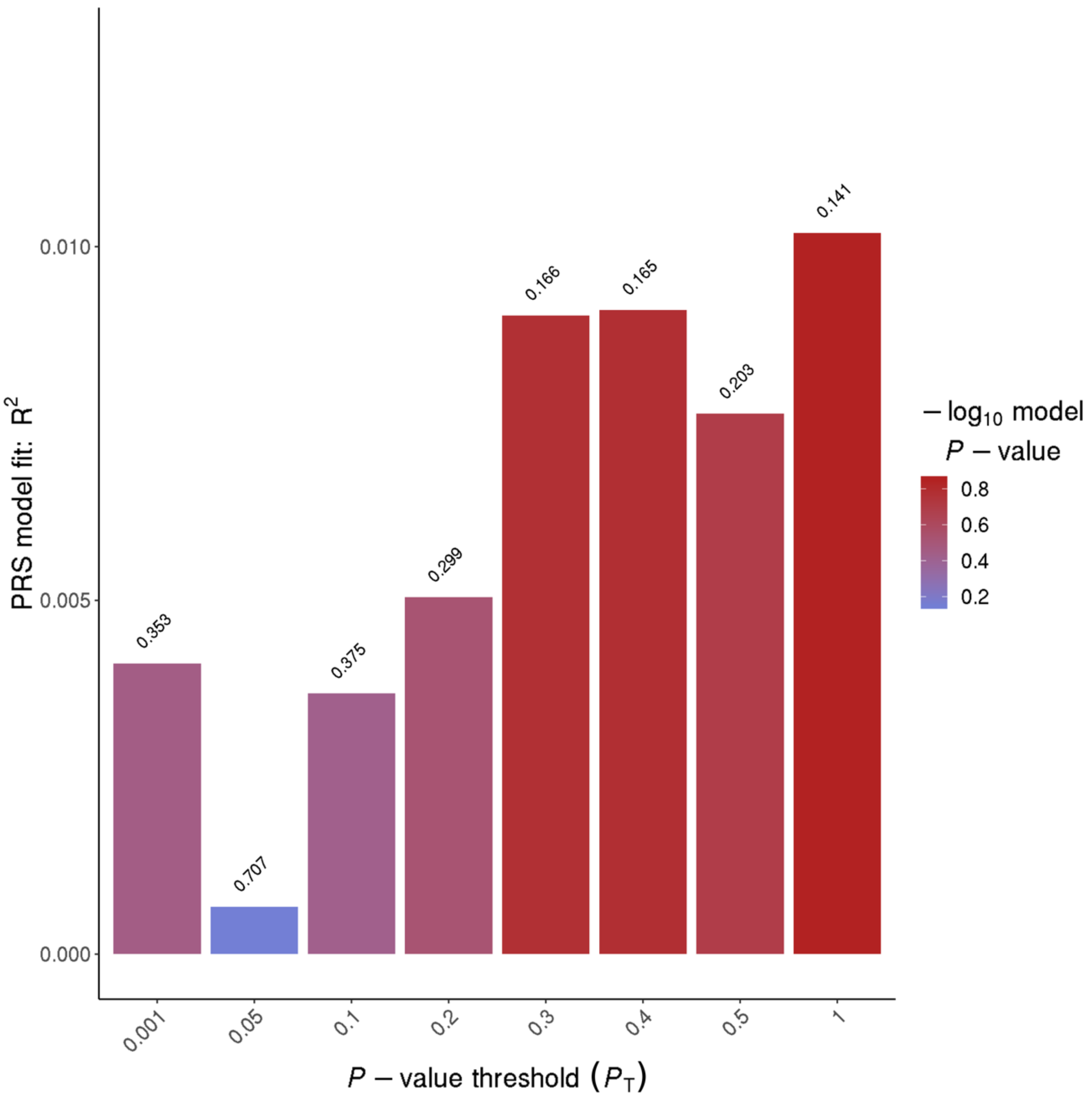


**eFigure 9: Bar plot from PRSice showing results at broad *P*-value thresholds for epilepsy PRS predicting status for 25 NAFE HS patients and 954 French-Canadian controls.**

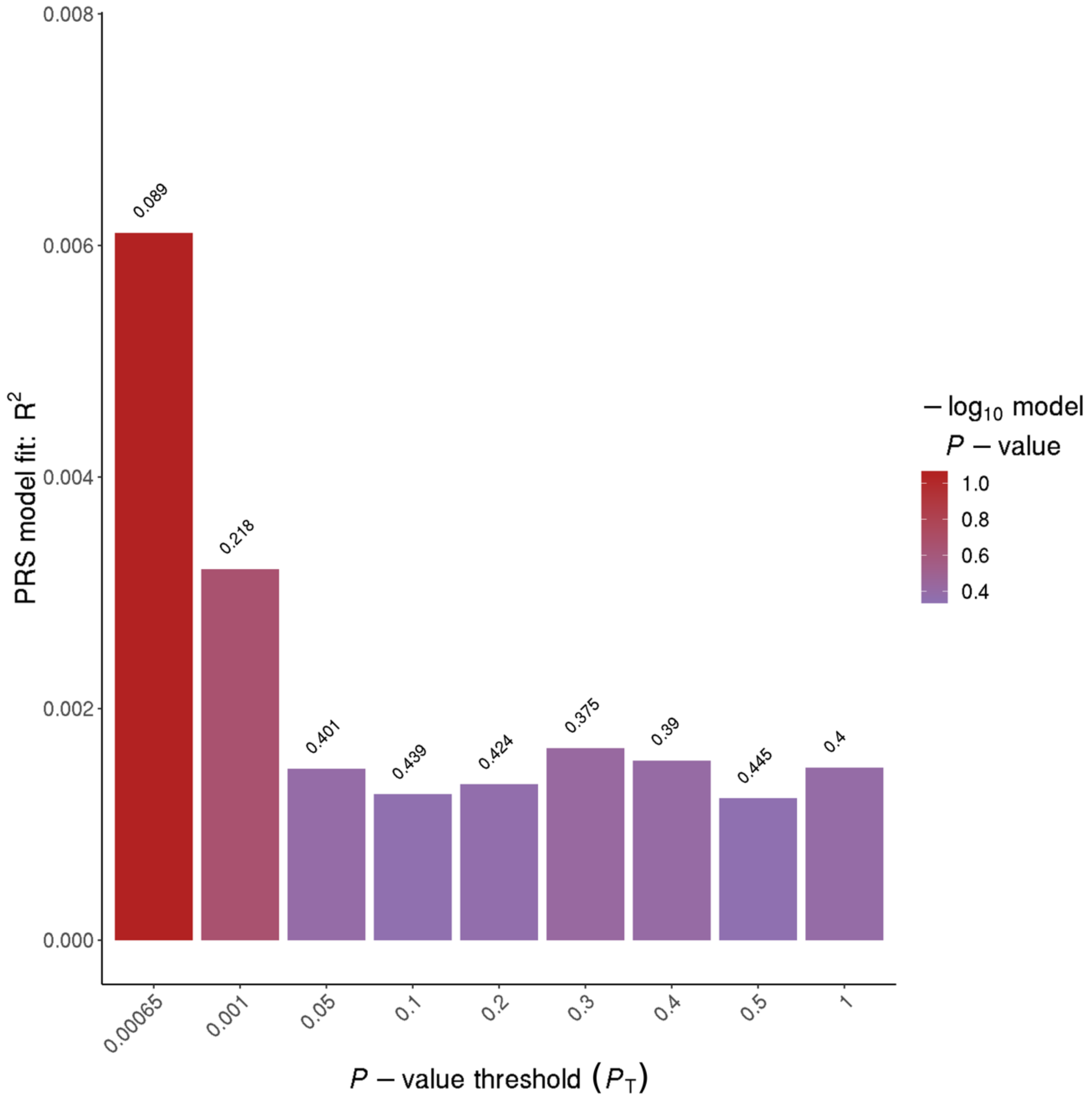


**eFigure 10: Bar plot from PRSice showing results at broad *P*-value thresholds for epilepsy PRS predicting status for 81 NAFE with no documented lesion patients and 954 French-Canadian controls.**

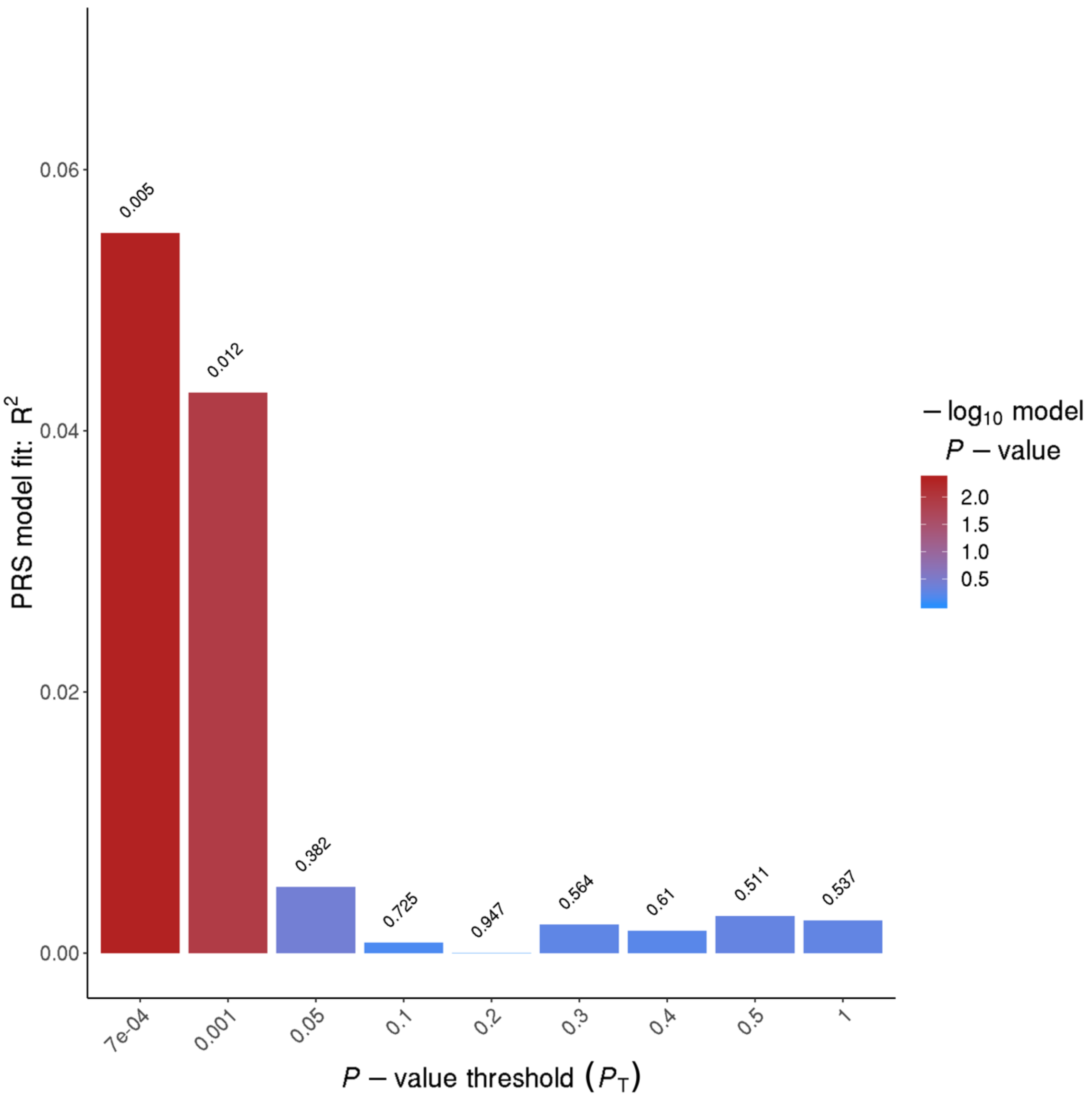


**eFigure 11: Bar plot from PRSice showing results at broad *P*-value thresholds for epilepsy PRS predicting status for 16 NAFE with documented lesion other than HS patients and 954 French-Canadian controls.**
